# Shark movements in the Revillagigedo Archipelago and connectivity with the Eastern Tropical Pacific

**DOI:** 10.1101/2020.03.02.972844

**Authors:** Frida Lara-Lizardi, Mauricio Hoyos-Padilla, Alex Hearn, A. Peter Klimley, Felipe Galván-Magaña, Randall Arauz, Sandra Bessudo, Eleazar Castro, Eric Clua, Eduardo Espinoza, Chris Fischer, César Peñaherrera-Palma, Todd Steiner, James T. Ketchum

## Abstract

Long-distance movements of sharks within and between islands pose substantial challenges for resource managers working with highly migratory species. When no-take zones do not cover the critical areas that sharks use as part of their lifecycle, exposure to fishing activities can be significant. Shark movements between the Marine Protected Areas (MPAs) of the Eastern Tropical Pacific (ETP) have been studied for several years, however little is known about the strength of connectivity between these islands. We analyzed the extensive MigraMar ultrasonic telemetry dataset to assess how Galapagos sharks (*Carcharhinus galapagensis*) and silky sharks (*Carcharhinus falciformis*) use different islands as stepping-stones during their migrations within the Revillagigedo National Park and other ETP islands. Of the 66 sharks monitored, 63.5% moved within the same island, 25.4% between two islands or more and only 10.1% across different MPAs. A *C. falciformis* tagged in Roca Partida Island, Revillagigedo, travelled to Clipperton Atoll and another one tagged in Darwin Island travelled to the atoll on two different years. The largest movement of *C. galapagensis* was accomplished by a shark tagged at Socorro Island, Revillagigedo, later detected at Clipperton and finally recorded in Darwin Island, Galapagos. This last path was in fact, one of the longest movements ever recorded for the species. Although long-distance dispersion was not common, our results highlight the need for co-operation between different countries to ensure adequate protection for sharks in the form of swimways and other conservation tools in the ETP.

## Introduction

Mobile top predators aggregate at specific sites or central locations near islands and seamounts, which they use for foraging, reproduction, thermoregulation and refuging, known as biological hotspots (1–3). Hence, movements of sharks and other marine predators need to be understood to: (i) assess how biological hotspots are inter-connected (2); (ii) define potential corridors of movement (4); and (iii) establish MPAs to provide more effective management actions for migratory species (5). Functional and physical links between different habitats, defined here as connectivity, are fundamental to maintain the biodiversity and resilience of an ecosystem (6). Knowledge of movement pathways in an area may help to: (i) inform management plans to maintain or restore connectivity (7); (ii) improve the design and effectiveness of MPAs; and (iii) define the functional role of a wide range of predators in marine ecosystems (8).

Many MPAs have been designed around oceanic islands to protect marine coastal and pelagic species such as sharks. Among notable MPAs in the ETP are the Galapagos Marine Reserve (138,000 km^2^), Malpelo Island Flora and Fauna Sanctuary (27,000 km^2^), and Cocos Island National Park (1,997 km^2^; 9). More recently, the territorial waters of Clipperton Atoll, France (8.9 km^2^ including the lagoon) was proposed as an MPA on 2016, however it is still unprotected (10). These areas not only contribute to the protection of species with high ecological value, but they also provide habitat for endangered species and are of paramount cultural value: all MPAs above mentioned have been designated as United Nations (UNESCO) World Natural Heritage Sites (11). UNESCO first recognized Cocos Island National Park in 1997, then the Galapagos Marine Reserve in 2001, Coiba National Park in 2005, Malpelo Flora and Fauna Sanctuary in 2006, and Revillagigedo Archipelago in 2016 (11). This last one is a reference of how large scale marine protected areas (LSMPAs) can be used to protect highly migratory species. In 2017, the MPA was expanded from 6,366 km^2^ to 148,000 km^2^ based on the inter-insular connectivity and the large-scale movements of different shark species (12), creating the largest no-take zone in North America (13).

Current literature shows that inter-island movements of sharks in the ETP are common. Evidence suggests that sharks may use islands as ‟stepping stones” for long distance oceanic dispersal (2, 4). For example, scalloped hammerhead sharks (*Sphyrna lewini)* at Wolf and Darwin islands in the Galapagos moved over 100 km to Roca Redonda and Seymour Norte within the marine reserve, and others made longer-distance movements across the ETP to other isolated islands, such as Cocos and Malpelo islands (4).

These movements in and out of MPAs imply that this species is vulnerable to both domestic fisheries within the Exclusive Economic Zones (EEZs) and multinational fisheries in the high seas (14, 15). Regular movements across MPA boundaries highlight the need for cooperation between jurisdictions to ensure sharks receive enough protection throughout their migrations (3). This need includes regulations focused on the habitats in each jurisdiction where individuals spend time, as well as movement corridors (16).

A recent initiative calls for the creation of *MigraVías* in Spanish (www.migramar.org), which are a set of connectivity conservation projects that create marine links and corridors between protected areas and other habitat patches such as seamounts, which increase the mobility and range of many species and allow them to move through the marine landscape so that genetic flow and diversity is maintained (17). By linking populations throughout the marine seascape, there is less chance of extinction and greater support for species richness and resilience of populations to climate change. The MigraVías are based on scientific evidence of organisms moving between different patches of habitat and work by increasing connectivity between these patches through conservation measures that allow them to move safely along marine corridors. Scientists and national governments are currently working together to create the first two in the Eastern Pacific: 1) the Coco-Galapagos MigraVías between Costa Rica and Ecuador, and 2) the Coiba-Malpelo MigraVías between Panama and Colombia (17).

The definition of the extent and occurrence of long-range movements and population connectivity are necessary for a full understanding of the ecology of a species and hence for designing effective conservation action (18). By assessing movement frequency, Network Analysis (NA) can be used to identify important movement corridors between core habitats of a species (6, 19). NA provides a new insight into the connectivity of specific habitats and the animals moving between them. It also proves valuable in revealing important information on distinct spatial and temporal changes in animal movements (6, 19). For example, an area with a high degree of centrality would suggest strong site fidelity by wide-ranging animals, hence the animals may return to the same location from many different areas (6).

Movements and residency patterns of key marine animals are still poorly understood, particularly within and between insular locations. In this study we describe the connectivity of Galapagos and silky sharks within and between insular sites in the Revillagigedo Archipelago and the ETP. The differences in the dispersal ranges of each species were detailed to identify the most important stepping-stones and movement MV in the region.

## Materials and methods

### Ethics statement

The work carried out in this study was done so in accordance with the following research permits (resolutions) from the Mexican Fisheries Commission (CONAPESCA) and the Revillagigedo National Park Authorities: SGPA/DGVS/06798; DRPBCPN.APFFCSL-REBIARRE.-067/2011; F00.1.DRPBCPN.-00405/0216; PPF/DGOPA-134/15; PPF/DGOPA-027/14; DGOPA.03624/240413; DGOPA.06668.150612.1691; F00.DFPBCPN.000211; DGOPA.10695.191110.-5322; DGOPA.042449.270409.-1151; SGPA/DGVS/06798; DRPBCPN.APFFCSL-REBIARRE.-067/2011; F00.1.DRPBCPN.-00405/0216; PPF/DGOPA-134/15; PPF/DGOPA-027/14; DGOPA.03624/240413; DGOPA.06668.150612.1691; F00.DFPBCPN.000211; DGOPA.10695.191110.-5322; DGOPA.042449.270409.-1151. Research methods for this study were approved by the Institutional Animal Care and Use Committee Protocol #16022, issued to the three co-authors based at the University of California, Davis at the time of the study.

### Species of interest

The silky shark is a globally distributed (40°N and 40°S) and highly migratory species (20, 21). It is found from the surface to depths of >200 m (22). Based on carbon (δ13 C) and nitrogen (δ15 N) isotope analysis, it was found that the species feeds in the open ocean, consuming oceanic pelagic prey (23, 24), normally at night or in the early morning (25, 26). It consumes squid, such as *Dosidicus gigas* during its vertical migration to the surface at night (27).

The Galapagos shark has a similar geographical (39°N-33°S) and depth distribution (from the surface to 180 m, but mostly <80 m), yet it is highly associated to seamounts, oceanic islands and continental shelf environments (28). Galapagos sharks feed primarily on demersal teleosts (29), but it can also consume cephalopods, elasmobranchs, crustaceans, small marine mammals (e.g. sea lions), and even other elasmobranch species (29).

### Study Area

The main study area is the Revillagigedo Archipelago (18°49′N 112°46′W), a group of four volcanic islands, 240 miles southwest of Cabo San Lucas, Mexico. The three eastern islands, San Benedicto, Socorro, and Roca Partida, called the inner islands, are relatively close to each other. Clarion is roughly 200 km to the west, and it is called the outer island. Socorro is the largest, covering an area of 132 km^2^. Clarion Island, the westernmost of the Archipelago, is the second largest of the islands, with an extension of 19.7 km^2^ in area (30).

We also studied the movements to and from other MPAs in the ETP. Clipperton Atoll (10°17’N 109°13’W) is positioned at the edge of the Eastern Pacific Barrier. This is the only coral atoll in the eastern Pacific which lies about 965 km from mainland Mexico. The 50-m isobath is ∼ 500 m from the reef front of this 3.7 km^2^ coral circle (30). Cocos Island (5°31’N 87°04’W) is located more than 500 km from mainland Costa Rica. It is the only point above sea level on the Cocos Ridge, which originates in the Galapagos Spreading Center. The 24 km^2^ island is surrounded by an insular platform that deepens to around 180 m, with an area of about 300 km^2^, then drops to several thousand meters deep (31). Malpelo Island (3°58′N and 81°37′W) is located 490 km from the Colombian Pacific coast. The 1.2 km^2^ island is surrounded by eleven pinnacles and its highest point is 300 m above sea level (32). The Galapagos Archipelago (0°40’S 90°33’W) is located 1,000 km from the coast of continental Ecuador. The archipelago is made up of 13 major islands and over 100 islets and emergent rocks, along with an unknown number of shallow and deep seamounts (33). The five ETP marine reserves (Revillagigedo, Clipperton, Cocos, Malpelo and Galapagos) are characterized by their complex oceanography, high diversity and abundance of pelagic species with high economical value for fisheries and tourism (34).

### Ultrasonic tag detection

Sixty-six sharks (32 *C. falciformis* and 34 *C. galapagensis*) were fitted with ultrasonic tags (Vemco, Ltd., Halifax, V16, frequency, 69 kHz, power 4-5H, life 1800 to 3650 days) during cruises to those five insular systems from 2006 to 2016. Tags emit a coded signal at 69 kHz with a random delay of 60–180 s to avoid successive signal collisions between two or more tags. Tags were fitted externally on sharks by scuba and free diving using pole spears or spearguns, inserting a stainless-steel barb into the dorsal musculature at the base of the dorsal fin. Other tags were implanted in the peritoneal cavity of sharks caught using hook and line. The gender (presence of claspers or not), maturity stage (neonate, juvenile, sub-adult, adult) and total length (estimated by free divers or measured for sharks which were captured) were recorded for all sharks when possible. Due to constant hurricanes in Revillagigedo during the wet season, tagging expeditions through the whole year were not possible. However, both species were tagged at the beginning, during and at the end of the dry season, expecting to reduce the bias for this study.

The Pelagios Kakunja (www.pelagioskakunja.org) and MigraMar Ultrasonic Receiver Networks (http://www.migramar.org/) registered the signals emitted by the ultrasonic tags. The receivers (Vemco Ltd., Halifax, VR2 and VR2W) were located in all oceanic MPAs across the ETP, spanning a straight-line distance of 4000 km from Revillagigedo to the Galapagos. The arrays were deployed between 2006 to 2012 at the following sites: Revillagigedo (Roca Partida, Clarion, Socorro and San Benedicto islands), Clipperton Atoll, Cocos Island, Malpelo Island and Galapagos Archipelago (Darwin, Wolf, Santa Cruz, Isabela, San Cristobal; Fig 1). Receivers were affixed with heavy-duty cable ties to a mooring line with chain or cable to attach to a bottom anchor and a buoy for flotation. Range tests of the ultrasonic receivers were performed for all the study areas, varying between 200 to 300 m (2, 35).

**Fig 1.**
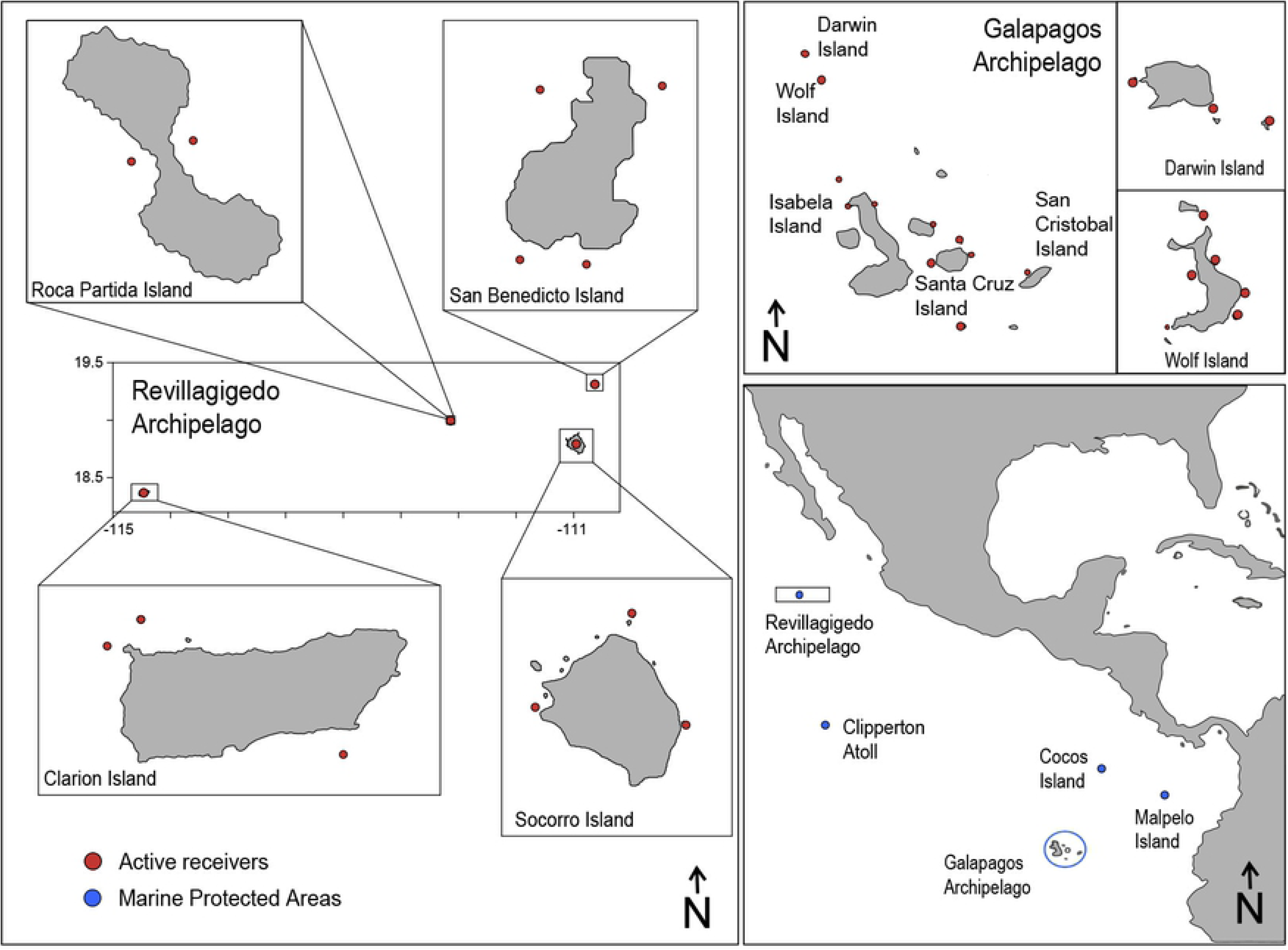
Map indicating the location of acoustic receivers used to monitor shark movements in the Revillagigedo National Park, the Galapagos National Park and the marine reserves in the Eastern Tropical Pacific.

### Data Analysis

#### Seasonality and distribution patterns

To describe the seasonality and residency of sharks in the Revillagigedo Archipelago through several years, we created point plots, which show the daily detections of each individual (y axis: Tag. ID) and the monitoring period (2010-2018). Then, we calculated the total number detections per day compared to the average Sea Surface Temperature, which was downloaded and filtered from the satellite Aqua MODIS from ERDAP NOAA web server (https://coastwatch.pfeg.noaa.gov/erddap/griddap; [37]). To compare the day/night distribution behavior of both species, we used circular plots, that summed the number of detections per hour, and the color shows a standard dial pattern (daytime: 6:00 – 18:00 h and nighttime: 18:00 – 6:00 h). We compared the distributions using the Rao test in R.

The ATT library (37) was used to calculate the Center of Activity (COA) and heatmaps. The following libraries of R were used to create maps: ggmap, osmdata and wesanderson (for color palette). A density function (Kernel Density Estimation) was plotted using the latitude and longitude of the Center of Activity database. We compared the movement activity between species and seasons: wet (June-November) and dry (December-May) in the Revillagigedo Archipelago.

#### Inter-island movements

To determine the proportion of individuals of *C. galapagensis* and *C. falciformis* that showed inter-island movements in and out of the Revillagigedo National Park, we counted the number of islands where each shark was recorded. Then, we plotted the detections over time, using examples of the different types of inter-island behaviors. To evaluate the dispersal range of each species (38) we measured the straight-line distances between acoustic receivers using the library geosphere in R.2.3.1 (R Core Team, 2017). We performed frequency histograms for the distance of each movement and compared the results from each species.

To describe the movement behavior of each species along the ETP we used NA using the igraph 1.2 package (39) available in the R programing language (R Core Team, 2017). The NA describes the local and global structure of networks constructed from pairwise interactions of connected elements in a graphic format node linked by one or a series of edges (6). In our analysis, each node represented the physical location of the acoustic receivers (hereafter sites). Edges were equally variable and represented the mobility of organisms between nodes. Each shark tagged represented a unique observation of the network. Several quantitative metrics were calculated from the interconnected network to describe the local and global network structure (19): (i) number of edges, (ii) number of vertices, (iii) degree of centrality and (iv) density. The density defined as the proportion of edges actually present in the network among all possible edges in the data (19). The degree of centrality defined as the overall level of connectedness within the network.

The NA was based on movements between receiver locations, where the size of the node represented the degree centrality (19). To determine the relative importance of each node within the marine reserves, we calculated the eigenvalues, defining the centrality of each node as a proportion to the sum of the centralities of those nodes to the ones which are connected. In general, nodes with high eigenvector centralities are those which are connected to many other nodes that are, in turn, connected to many others (19).

## Results

### Seasonality and residency over the years

Sixty-six sharks (34 *C. falciformis* and 32 *C. galapagensis*) were tagged and monitored since 2010 (Table 1; Fig 2). Some sharks were observed for over a period of five years. We plotted the wet (stormy) and dry seasons to observe the relationship of shark detections with changes in water temperature. During the stormy season, the number of movements were reduced, which also shows drastic changes in the sea surface temperature of almost 10°C in a single day (Fig 3). Although both species showed similar distribution patterns, when we compared their records according to the dial cycles, *C. falciformis* showed higher number of detections during daytime. Whereas, *C. galapagensis* was recorded mainly during nighttime, the Rao’s test showed a significant difference (p<0.001). This means the detections during the day were not random and that some hours are more important than others (Fig 4).

**Fig 2.**
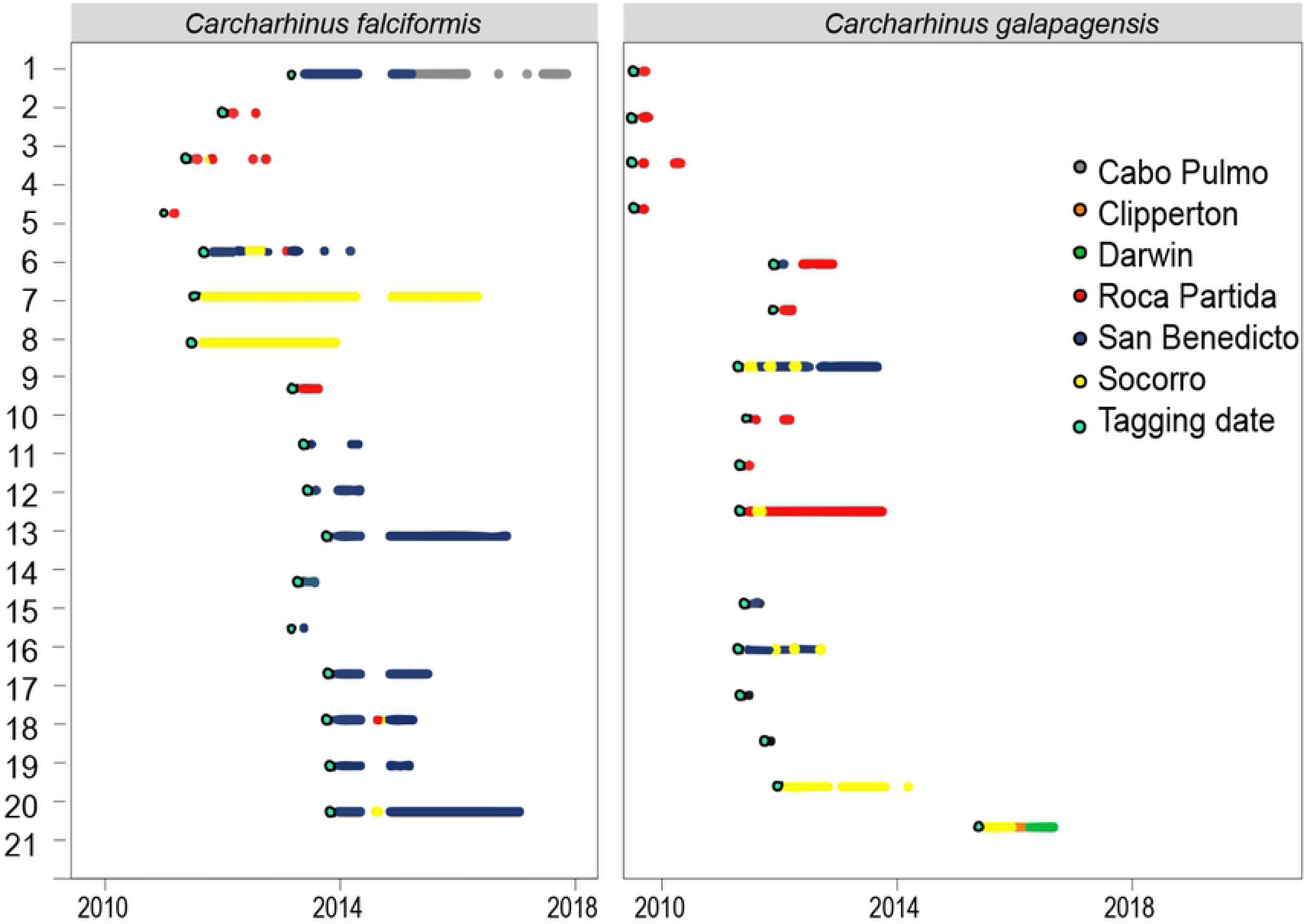
Chronology of detections of *C. falciformis* and *C. galapagensis* over the last eight years in Revillagigedo National Park. Islands are indicated by colors and initial tagging by a light blue dot.

**Fig 3.**
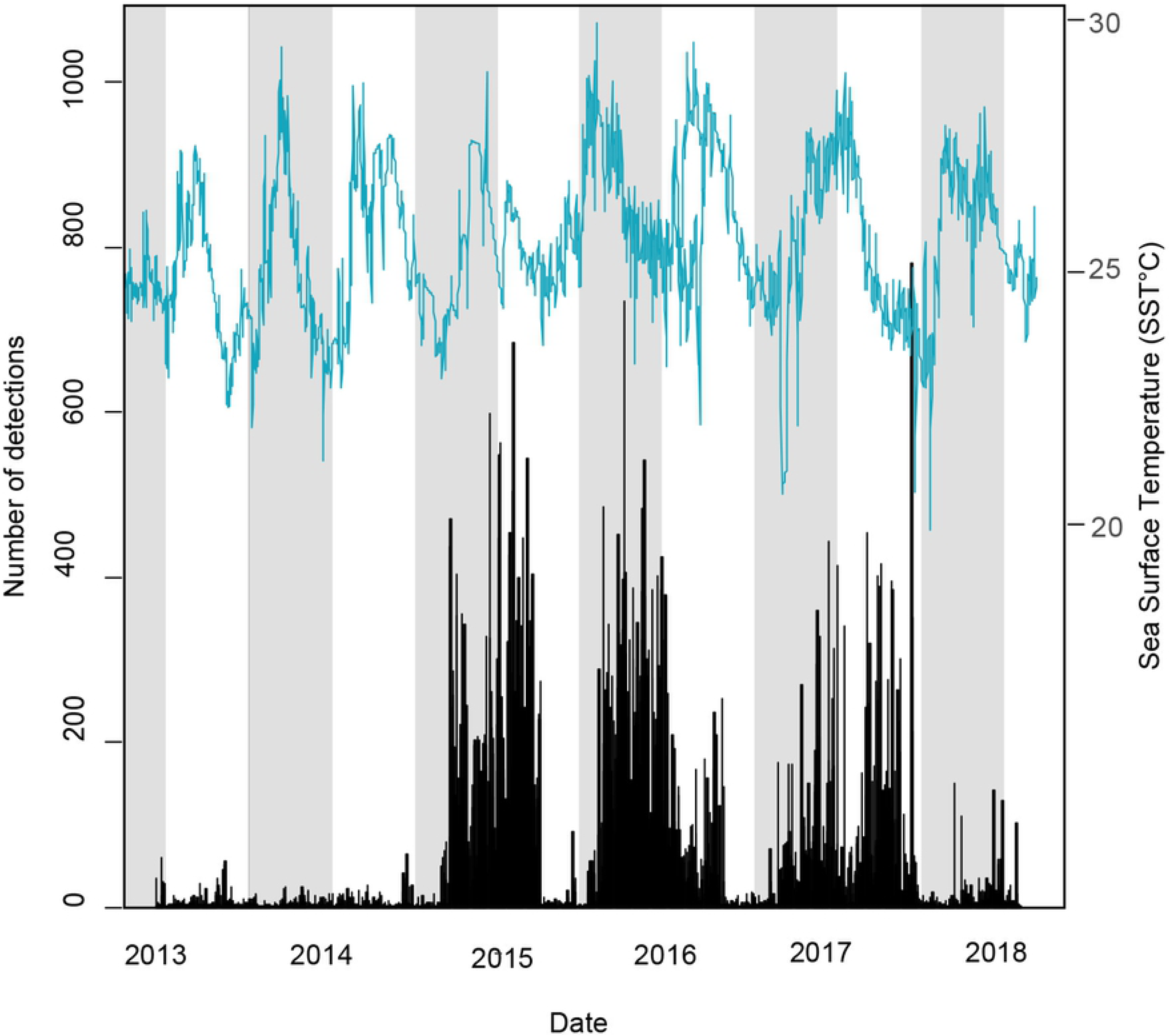
Daily detections of *C. falciformis* and *C. galapagensis* according to the sea surface temperature (SST) during the last five years of monitoring in Revillagigedo National Park. The white bars indicate the wet season and grey bars show the dry season.

**Fig 4.**
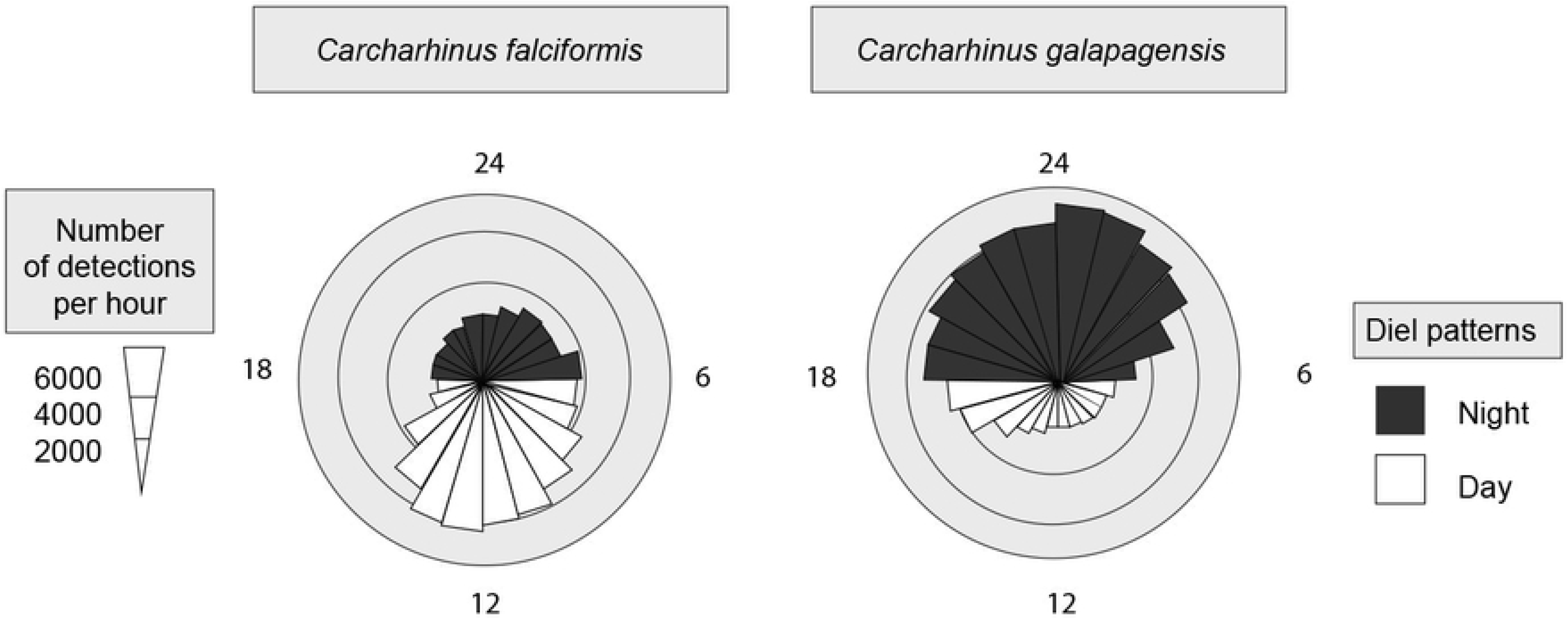
Diel presence of *C. falciformis* and *C. galapagensis* at receiver locations in the Revillagigedo National Park.

**Table 1.** Tagging information of the sharks monitored in Revillagigedo National Park (2010-2017).

|  | *C. falciformis* | *C. galapagensis* |
| --- | --- | --- |
| **Females** | 30 | 21 |
| **Males** | 1 | 5 |
| **Unknown** | 1 | 8 |
| **Total** | 32 | 34 |

### Center of activity (COA) and seasonal interisland movements

The activity maps were based on acoustic detections and their COA showing the sites with the highest number of detections. Comparing the seasons, there is a significant higher activity density that can be observed in the wet season compared to the dry season for *C. falciformis* (p<0.05; Fig 5). There was an increase in the number of movements and sites visited by the silky and Galapagos sharks monitored in Revillagigedo in the wet season, compared to the dry season (Fig 6).

**Fig 5.**
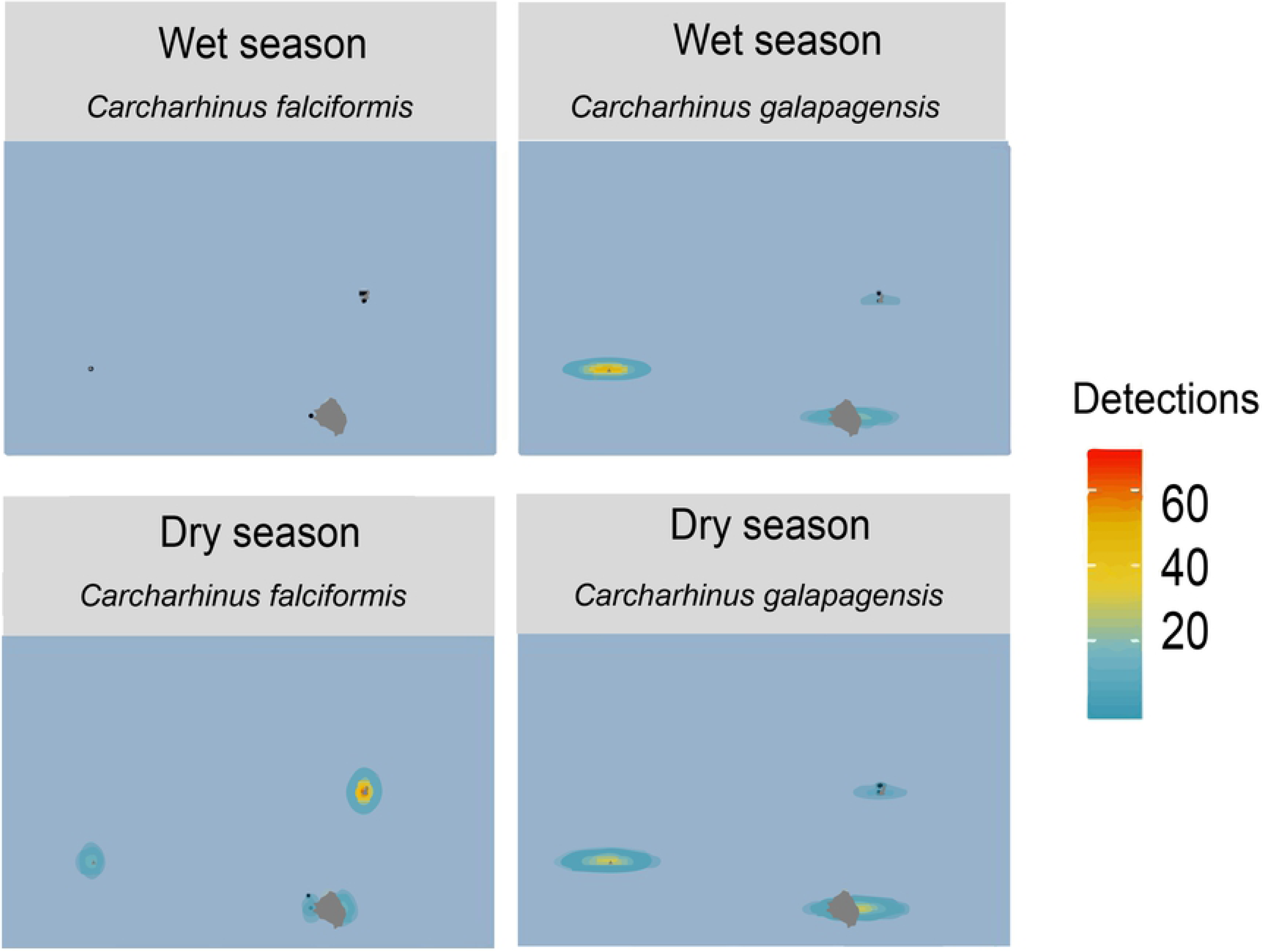
Density plots showig of the movement of silky and Galapagos shark, based on acoustic detections and their center of activity (COA) in Revillagigedo National Park during the dry (top) and wet season (bottom), where red indicates the sites with the highest number of detections and green sites indicate the lowest activity.

**Fig 6.**
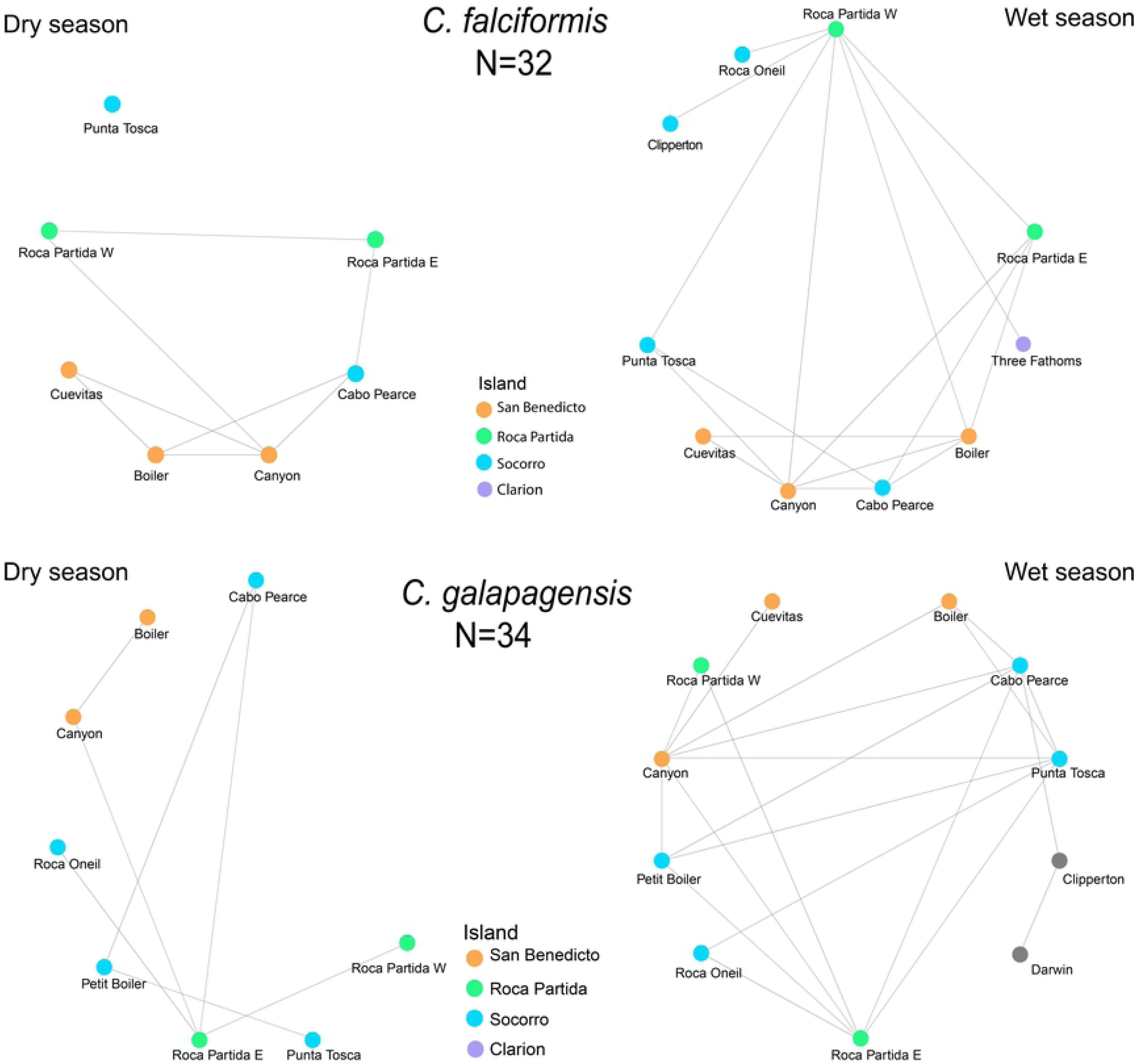
Network analysis of *C. falciformis* (top) and *C. galapagensis* (bottom) monitored in Revillagigedo National Park.

### Daily and inter-island movements

Of the overall detections analyzed, 63.5% of individuals undertook movements within the same island, 25.4% between islands and 10.1% across MPAs (Fig 7 and 8). For example, the Galapagos sharks tagged at Socorro spent only a few days at a time at the first island, while they stayed longer at Roca Partida and San Benedicto. Another shark tagged at San Benedicto in early January, visited Socorro Island for two months, from the middle of March to the middle of May, before returning to San Benedicto. It made another brief visit to Socorro later that year. The silky sharks tagged at the Revillagigedo Archipelago were resident for a period of one month to four months, others visited for a day or two before leaving the islands.

**Fig 7.**
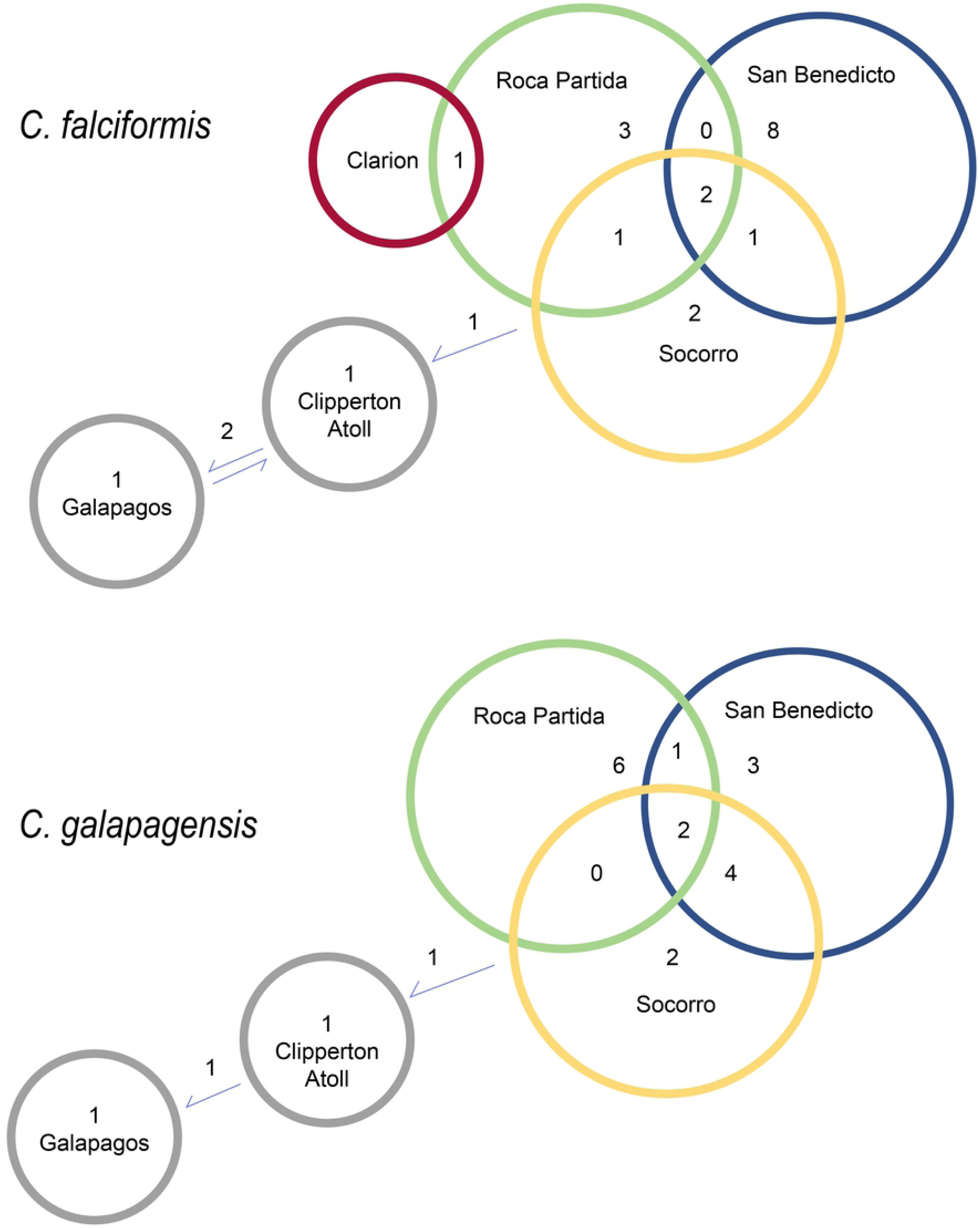
Number of individuals of *C. falciformis* (left) and *C. galapagensis* (right) that showed inter-island movements in and out of the Revillagigedo National Park.

**Fig 8.**
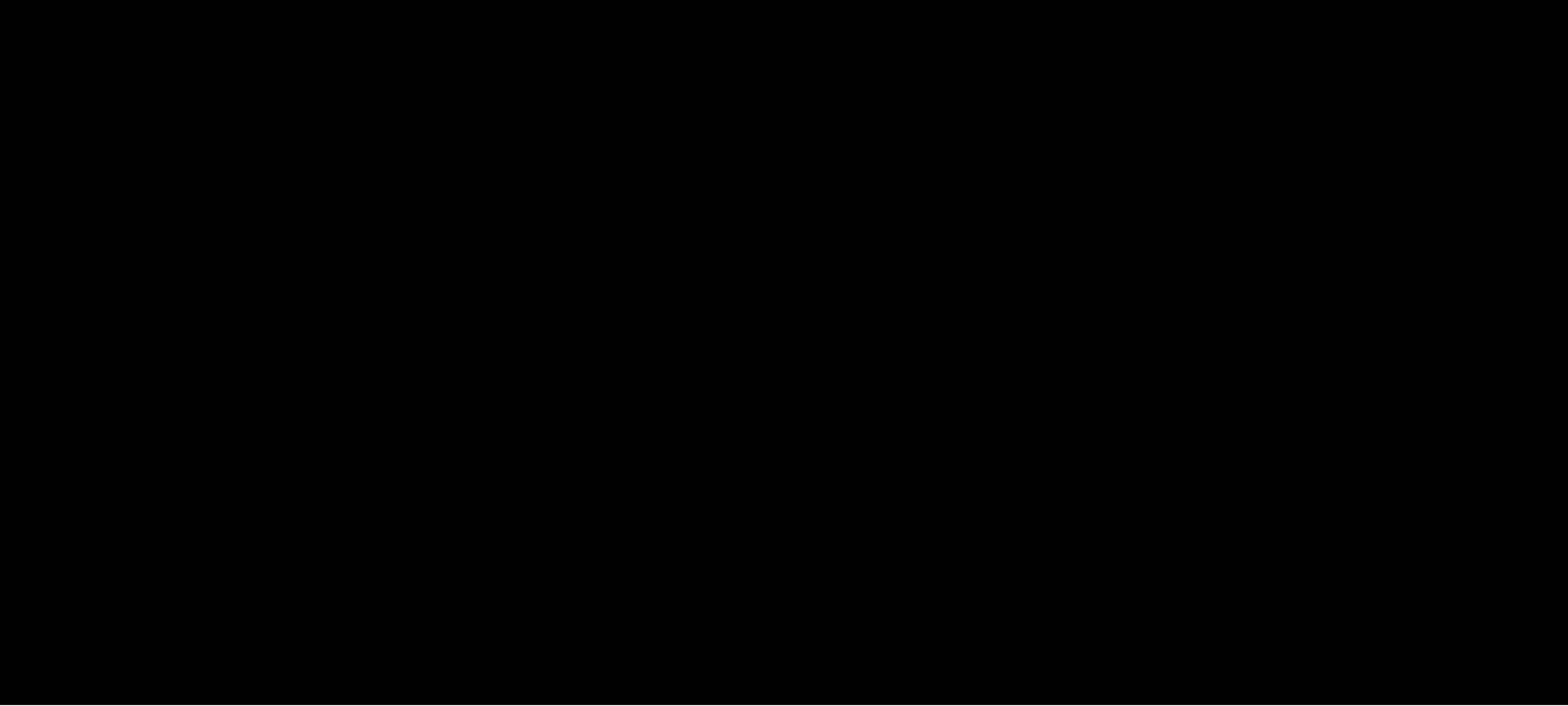
Inter-Island movements of *C. falciformis* and *C. galapagensis* recorded in the Revillagigedo National Park. The red circle represents the previous no-take, the shaded gray circles shows the proposed no-take area and the black polygon shows the current extent of the Revillagigedo National Park.

### Movements between MPAs

Despite these large-scale records, 90% of the movements of both species were observed in a range of < 50 km (Fig 9), showing high site fidelity to the tagging site. A female silky shark of 163 cm total length (TL), tagged in Roca Partida Island, Revillagigedo, travelled to Clipperton Atoll (965 km to the south). Another female silky shark of 187 cm TL, tagged in the anchorage at Wolf Island, Galapagos, travelled to Clipperton Atoll and back again (2,200 km to the north) in two different years (Fig 10). In contrast, the largest movement of *C. galapagensis* was accomplished by a female with a total length of 180 cm TL, tagged on February 2016 at Socorro Island, Revillagigedo, later detected at Clipperton Atoll (960 km south of the tagging site) and then in March 2017 again recorded in Darwin Island, Galapagos (2,200 km to the south), which is one of the longest movements ever recorded for the species (Fig 11).

**Fig 9.**
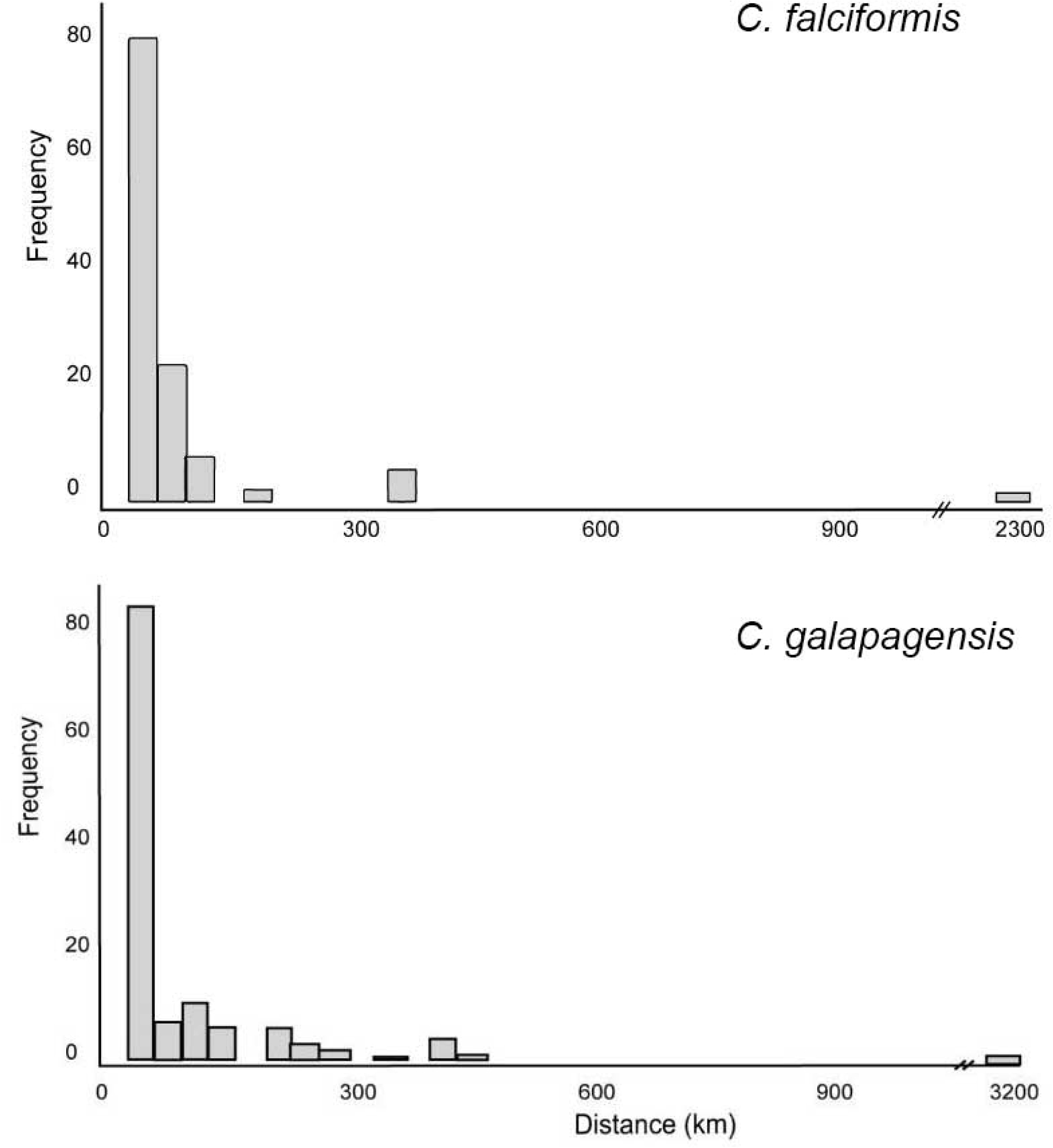
Frequency of sharks’ *C. falciformis* (on the top) and *C. galapagensis* (on the bottom) movements per distance (kilometers) of the Revillagigedo National Park.

**Fig 10.**
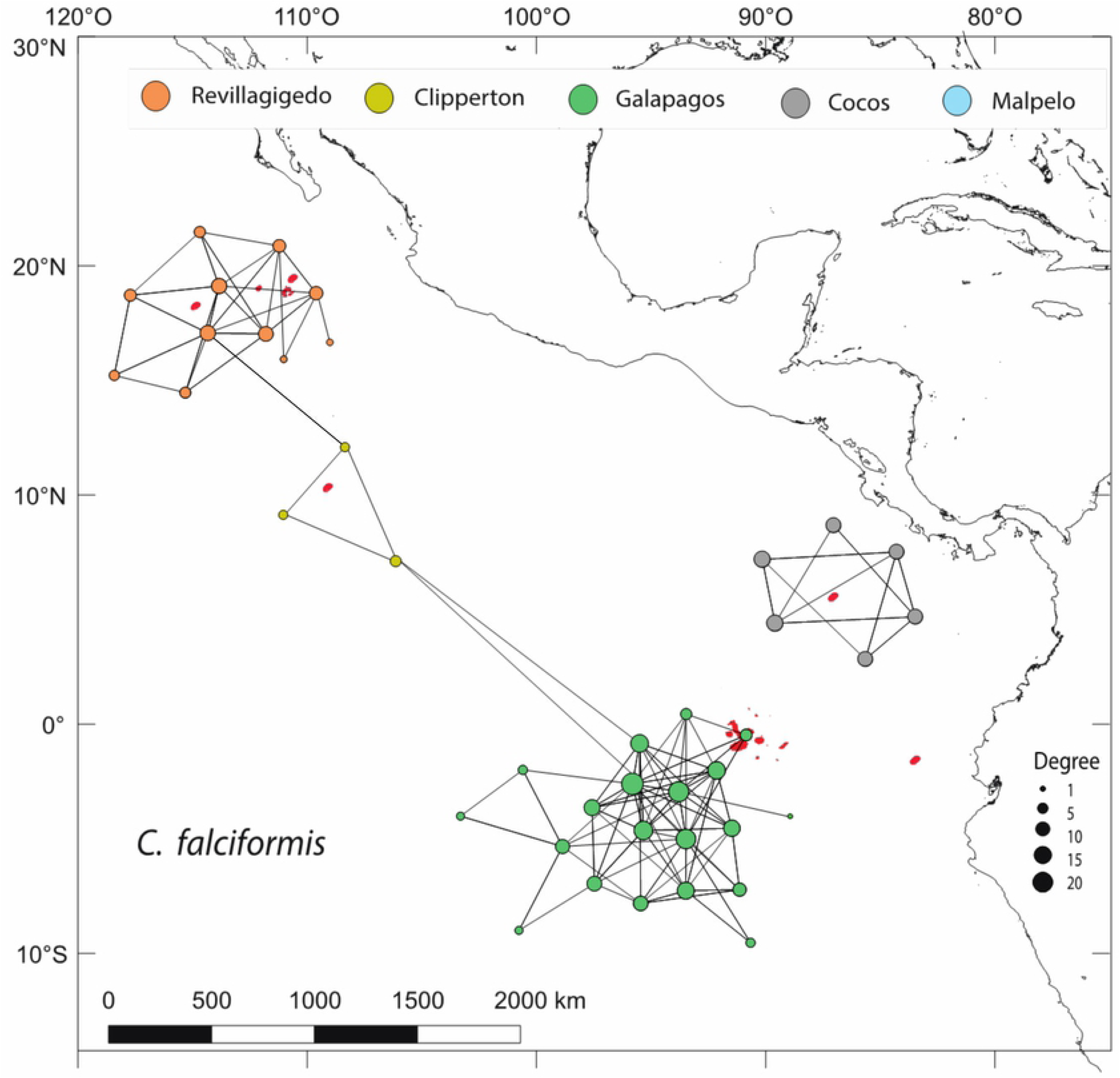
Network analysis of *C. falciformis* monitored in the Eastern Tropical Pacific. Circles represent the nodes and the arrows indicate the edges or movement paths. The size of the circles represents the degree, that is, the number of links for each receiver.

**Fig 11.**
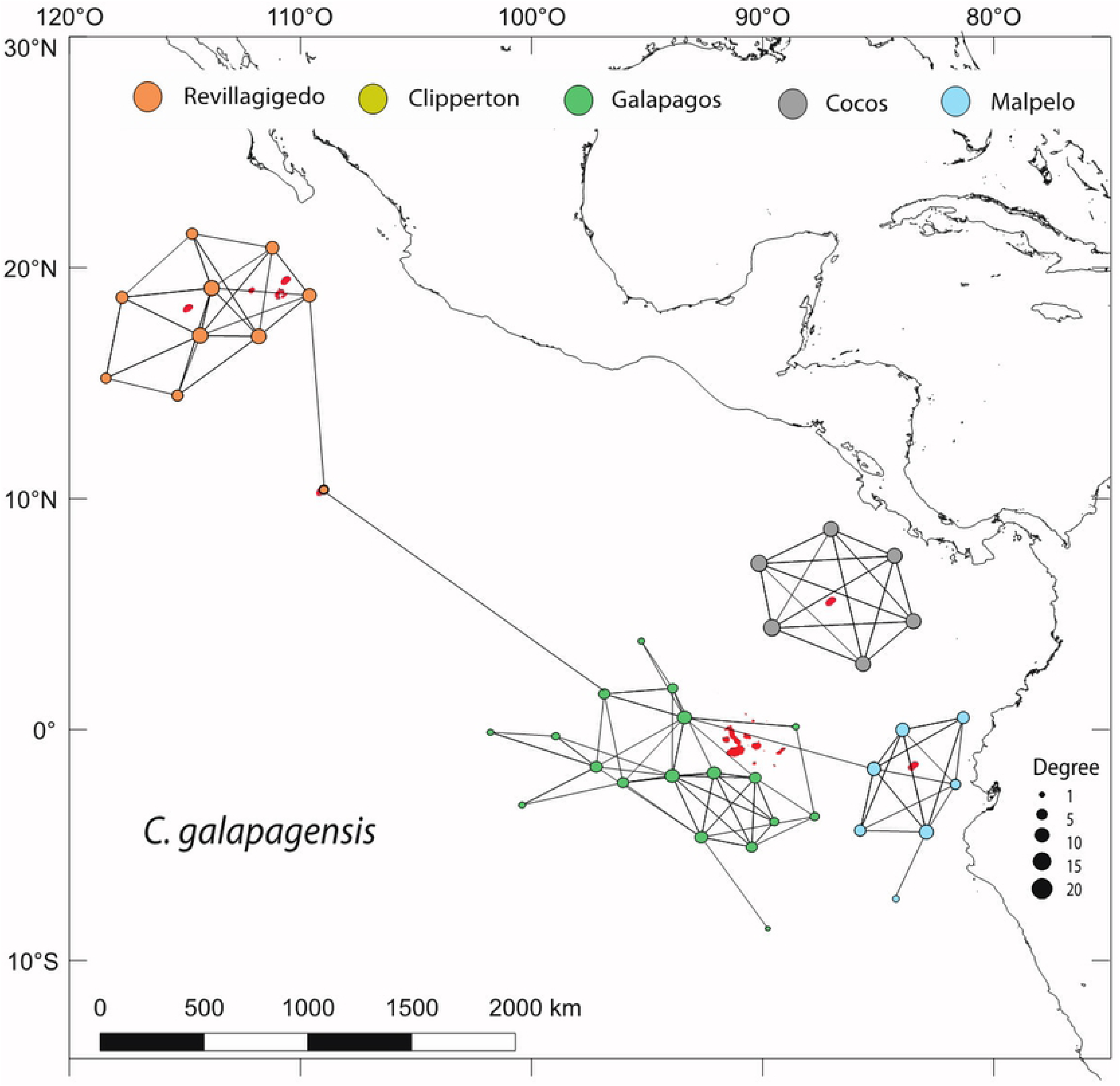
Network analysis of *C. galapagensis* monitored in the Eastern Tropical Pacific. Circles represent the nodes and the arrows indicate the edges or movement paths. The size of the circles represents the degree, that is, the number of links for each receiver.

### Network analysis and metrics

According to the NA metrics, *C. falciformis* has a more complex network with significantly higher values than *C. galapagensis* in terms of the number connections between the nodes or “edges” (X^2^= 44.714, df=1, p<0.5), the complexity of the network or “degree of centrality” (X^2^= 40.164, df=1, p<0.5) and the proportion of nodes used of the total options or “density” (X^2^= 14.238, df=1, p<0.5), whereas the number of recorded stations or “nodes” did not show a significant difference between the two species (X^2^= 0.10001, df=1, p= 0.75).

According to the NA, *C. falciformis* presented significantly higher values than *C. galapagensis* in terms of the number edges, the centralization and the density (p<0.5), whereas the number of nodes did not show a significant difference (p= 0.75). The eigenvector measured how well-linked a site was within the network; sites with a high eigenvector had high node strength and were connected to sites with similarly high node strength. The most important sites in terms of the eigenvalue varied according to the species, for *C. falciformis* the most important were Canyon (Revillagigedo), Darwin Anchorage (Galapagos) and Lobster (Cocos). Whereas,for *C. galapagensis*, the most important sites were Roca Partida (Revillagigedo), Nevera (Malpelo), Roca Elefante and Corales Norte (Galapagos).

## Discussion

According to this study, *C. falciformis* moved more frequently to nearby sites, but also showed high fidelity to their tagging location. Some sharks may gain more protection because of the location of their highest movement site and/or the distribution of management zones in the system (2, 40). Consequently, targeting specific sites based on prior knowledge and increasing the level of protection to include closely spaced habitats (20 km) may perform better for species like *C. falciformis* than having a single reserve.

During this study, one of the longest movements for *C. galapagensis* has been recorded. A subadult female of 181 cm TL was tagged in Socorro Island the 26^th^ of February 2016, then was detected 945 km south in the Clipperton Atoll for three months and finally it was detected in Darwin Island, Galapagos (2,300 km south). Therefore, this single individual showed a movement of at least 3,200 km south, passing by three marine reserves. In the same way, a female silky shark tagged in the anchorage at Wolf Island, Galapagos, travelled to Clipperton Atoll and back again (2,200 km to the north) in two different years, showing that this species present very long movement patterns as it was expected.

Shark populations are not homogenously distributed in different habitats of the ecosystem that can support a higher diversity and abundance (2, 5). Many shark species are known to aggregate on outer parts of reef slopes that are generally exposed to stronger current flow (3,4,41), where productive foraging grounds are present (40). Hence, currents probably shape the shark community and define spatial and temporal patterns of habitat use.

Hearn *et al.* (2) and Ketchum *et al.* (15) provided evidence to support this hypothesis by showing that specific areas around Wolf Island (Galapagos Marine Reserve), with stronger current flow, were generally ‘hotspots’ for hammerhead sharks and for other pelagic species, including Galapagos sharks. Based on the NA results, sharks are not just highly residential, but they also start long dispersal from these sites to other islands and marine reserves (more than 100 km).

We determined that these stepping-stones are sites where earlier studies have found high abundance of sharks. In Cocos, studies have shown that there are less sharks in sheltered bays, than in islands and seamounts (4, 42) For example, Nalesso *et al.* (4) found that Dos Amigos, Roca Sucia, and Alcyone are the sites with the highest abundance for the scalloped hammerhead, *S. lewini*. Manuelita also is important, but it varies according to the habitats within the site (2). In the Galapagos and Revillagigedo Archipelago, sharks seem to show a similar behavior as in Malpelo, with the largest aggregations found up-current in the side of the island where the current flows into (43). Ketchum *et al. interisland movements* (15) also mentioned that Darwin Island may be a stopover site for hammerheads that perform long-distance movements.

Few inter-island movements were observed within the marine reserves, and most of the movements were within 50 km. Previous studies have shown that female Galapagos sharks display high site-fidelity, while males are less resident (44). In general, they show movements of less than 100 km (44, 45). The longest travelled distance recorded for a Galapagos shark reached 2,859 km, a male that moved off Bermuda into the central Atlantic Ocean. We recorded a movement of *C. galapagensis* between Revillagigedo to Galapagos of 2,958 km, one of the longest movements reported in the ETP. Based on our results, the connectivity between the MPAs of the ETP relies on very few animals. A stronger connectivity is expected by increasing the tagging effort in the region. However, differences in receiver network deployment and acoustic coverage also affected the results. The analyses did not consider the distance between the receivers, therefore the probability of detecting more movements in short distances was expected. Heupel *et al.* (46) determined that for wide-ranging species, there is an under-estimation of the connectivity, because some individuals can appear to be absent from receiver locations for long periods while remaining within the general study area but outside the detection range of the receivers. As a critical finding, the long distances movements recorded in this study show the potential population connectivity within the ETP.

Pazmiño *et al.* (47) used a combination of mtDNA and diagnostic nuclear markers to properly assess the genetic connectivity of the Galapagos shark across the ETP and detect patterns of hybridization. The records of hybrids (Galapagos and dusky shark, *C. obscurus*) showed that these are migrating from the area of contact, the Revillagigedo Archipelago towards the Galapagos Islands using Clipperton Atoll as a stepping-stone. However, only 1% of the total sampled sharks showed this pattern.

Defining these movements between habitats is important to identify critical environments or corridors that may be important for population connectivity zonation (14, 48) and developing management strategies that ensure protection (49).

Both species can move between these two widely separated sites, but how often these movements occur is unclear – and the shedding of externally attached tags makes it likely that tags do not stay on long enough to get infrequent long-distance migrations. Furthermore, the detection of their presence at the two widely separated sites is based on receiver detections.

Clipperton Atoll is an area with unusual assemblages of both Indo-Pacific and Panamic flora and fauna (10), and it is possible that it is an important stepping-stone for connection between the two bioregions, Northern ETP (Revillagigedo and Gulf of California, e.g. 14) and Southern ETP (Malpelo, Cocos and Galapagos, e.g. 8).

It is becoming increasingly clear that some species can benefit from investments in local conservation measures nested within broader international efforts. However, Kinney et al. (50) established that nursery closures or size limits that protect only neonates and young juveniles are unlikely to fully promote population recovery, that is, effective management must involve protection for older age classes along with nursery-using life stages.

The ideal MPA design provides protection for all life stages of the species of concern, which is impractical for most shark species because they are wide ranging. According to the results, conservation within insular zones of the ETP region may have broad geographic benefits, because these reserves may be efficient protective zones, as long as they have a minimum size of 70,000 km^2^ established around each island, where persistence is highest and vulnerability is lowest (51).

In that perspective, the positive step forward based on the protection of the territorial waters of Clipperton Atoll should be continued with an extension of the protection area to reach this critical threshold of 70,000 km^2^ (e.g. 12). The observed movements between marine protected areas suggest that these species are vulnerable to domestic fisheries as well as multinational fisheries on the high seas, as these species are highly associated with commercial pelagic species such as, yellowfin tuna, *Thunnus albacares* (26). The preference of *C. galapagensis* to remain at or above 50 m depth makes the species much more vulnerable when moving offshore between reserves (29). Furthermore, even when not targeted, these sharks often comprise a high proportion of landings in line-based fisheries (52). For example, Kohin *et al.* (25) determined that silky sharks tagged in Costa Rica ranged into the EEZ of six countries and beyond into international waters. Increased protection of reefs and inter-reef habitats along the inner shelf may provide a greater conservation benefit. Definition of the extent and occurrence of long-range movement and population connectivity is necessary for a full understanding of the ecology of a species and hence for designing effective conservation action.

However, it has been recognized that the ETP region has a poor level of enforcement of laws. There is a low capacity to detect and intercept offenders, poor preparation for effective legal cases, difficulties in both administrative and judicial processes, and finally, obstacles which prevent sanctions from being imposed upon violators (9). Therefore, the use of new technologies and international agreements should be applied more often.

## Conclusions

Regular movement across state boundaries highlights the need for cooperation between countries to ensure that sharks receive enough protection throughout their migrations. This may include the need for regulations related to the habitats in each jurisdiction where individuals spend time, as well as movement corridors, such as the proposed swimways (MigraVías) in the ETP.

In Revillagigedo, the center of activity showed that sharks tend to be present during the wet months and they do move between sites during these months. The shark movement seasonality is related to current exposure, storms and temperature. According to the network analysis there are movements between MPAs but are not very common. However, they are very important in terms of conservation and management. Further research will elucidate the importance of the MigraVías showing how often biological corridors are utilized by migratory species.

## Acknowledgements

The authors would like to thank the International Community Foundation, Ocearch, Chris Fischer Productions and National Geographic for providing the initial funding to tag a large number of sharks and set up the acoustic receiver array in the Revillagigedo Archipelago. We are gratefiul to Club Cantamar, Fins Attached, Abismar, Sharks Mission France and Quino El Guardian for supporting our expeditions to the islands. We also thank the Alliances WWF-Telmex-Telcel and WWF-Fundación Carlos Slim for additional support for this project, as well as the Secretariat for Higher Education, Science, Technology and Innovation of the Ecuadorean Government; ICDF; Iris and Michael Smith; the Directorate of the Galapagos National Park. FGM thanks Instituto Politécnico Nacional (COFAA, EDI) for fellowships. This research was carried out under permits from the Secretaría de Agricultura, Ganadería, Desarrollo Rural, Pesca y Alimentación (DGOPA.06668.150612.1691) and Comisión Nacional de Áreas Naturales Protegidas (F00.DRPBCPN-APFFCSL.REBIARRE-102/13) of Mexico. We are also grateful to Secretaría del Medio Ambiente y Recursos Naturales and Dirección del Parque Nacional Revillagigedo for providing necessary permits to conduct research at the Revillagigedo National Park, a UNESCO World Heritage Site.

## Supporting information

S1 Fig. Number of sharks tagged by month in Revillagigedo National Park.

S2 Fig. Network metrics of *C. falciformis* and *C. galapagensis* of the ETP comparing the species.

